# ROS-dependent innate immune mechanisms control *Staphylococcus aureus* MRSA virulence in the *Drosophila* larval model

**DOI:** 10.1101/2020.10.02.323444

**Authors:** Elodie Ramond, Anne Jamet, Xiongqi Ding, Clémence Bouvier, Louison Lallemant, Xiangyan He, Laurence Arbibe, Mathieu Coureuil, Alain Charbit

## Abstract

Antibiotics multi-resistant *Staphylococcus aureus* strains constitute a major public health concern worldwide and are responsible of both healthcare- and community-associated infections. Here we have established a robust and simple *S. aureus* oral infection model, using *Drosophila melanogaster* larva, which allowed to follow *S. aureus* fate at the whole organism level as well as the host immune responses. Fluorescence microscopy and Light sheet 3D imaging revealed bacterial clustering at the posterior midgut that displays neutral pH. Our study demonstrates that *S. aureus* infection triggers host H_2_O_2_ production through Duox enzyme, consequently empowering antimicrobial peptides production through Toll pathway activation. We also show that catalase-mediated quenching of H_2_O_2_ not only enhances *S. aureus* survival but also minimizes host antimicrobial response, hence reducing bacterial clearance *in vivo.* Finally, we confirm the versatility of this model by demonstrating the colonization and host stimulation capacities of two other bacterial pathogens: *Salmonella* Typhimurium and *Shigella flexneri.* Overall, the drosophila larva may constitute a general model to follow *in vivo* host innate immune responses triggered upon infection with human pathogens.

## Introduction

*Staphylococcus aureus* is a facultative aerobic Gram-positive bacterium that behaves as a commensal microorganism (up to 30% of the healthy human population carries *S. aureus* through nasal, skin and intestinal colonization) or as a pathogen causing wide range of infections in humans, wild and companion animals (Matuszewska et al. 2020; Sivaraman et al. 2009; Parlet et al. 2019). The emergence of methicillin-resistant *S. aureus* (MRSA) clones that express numerous virulence factors including toxins and adhesins, increasing their toxicity and colonization capacities, is a major public health issue. Expression of these numerous virulence factors are correlated with severe symptoms among previously healthy colonized individuals (Lakhundi & Zhang 2018; Thurlow et al. 2012). *S. aureus* can also behave as an opportunistic pathogen in individuals with underlying diseases such as inflammatory bowel disease (*e. g*., Crohn’s disease) (Bettenworth et al. 2013). During infection, *S. aureus* must face host innate immunity *i.e.* phagocyte-mediated elimination via oxidative stress (by macrophages and neutrophils) and antimicrobial peptides secretion (DeLeo et al. 2009). *S. aureus* undergoes both endogenous oxidative stress (notably caused by incomplete aerobic respiration (Gaupp et al. 2012; Kohanski et al. 2007)) and exogenous host-induced oxidative stress aimed at killing the bacteria. At the site of infection, ROS are secreted in the extracellular space or in specialized phagocytic cells through Nox/Duox NADPH (Nicotinamide Adenine Dinucleotide PHosphate) oxidases. The Nox and Duox families are composed respectively of five Noxs (Nox1-Nox5) and two Duoxs (Duox1-Duox2) members. Nox1 and Duox2 are found in the gastrointestinal tract whereas Nox2 was identified in phagocytic cells (Bhattacharyya et al. 2014; Aviello & Knaus 2018). All other enzymes were identified in other tissues such as airway epithelium, kidneys, endothelial cells, *etc*… (Brown & Griendling 2009; Soodaeva et al. 2019). Nox enzymes catalyze superoxide production (O_2_^.−^) whereas Duox enzymes trigger hydrogen peroxide (H_2_O_2_). When phagocytosed by neutrophils or macrophages, *S. aureus* has to deal with the Nox2-dependent superoxide anion and their derivates (Buvelot et al. 2017). To neutralize ROS deleterious effects, *S. aureus* expresses multiple detoxifying enzymes including the expression of: i) the two superoxide dismutases SodA and SodM (*sodA* and *sodM* genes), which convert superoxide anions to H_2_O_2_ and O_2;_ ii) the H_2_O_2_ detoxifying enzyme catalase (*katA* gene); and iii) the alkyl hydroperoxide reductases (*ahpC* and *ahpF* genes), that neutralize H_2_O_2_ and alkyl hydroperoxides (Beavers & Skaar 2016). Currently, little is known about *S. aureus* adaptive mechanisms to oxidative stress.

Although most prior studies relied on the mouse model, mechanistic and genetic analyses can be performed with powerful alternative animal models such as *Drosophila melanogaster* (fly) or *Danio rerio* (zebrafish). Notably, use of zebrafish for gastrointestinal tract studies shows several disadvantages for intestinal infection researches as pH variations are distinct from mammalian intestine (pH remains around 7.5) (Nalbant et al. 1999; Brugman 2016). Moreover, *Drosophila* microbiota, corresponding to *Acetobacter* and *Lactobacillus* genera (Wong et al. 2011; Fink et al. 2013), are closer to human microbiota, in comparison to the zebrafish that is mainly colonized by the *γ-Proteobacteria* class, and specifically by the genera *Aeromonas* (Stephens et al. 2016). Of particular interest, flies and human share many similarities regarding physiological and anatomical aspects, especially about intestinal organ (Mistry et al. 2016). When infected, flies have the ability to generate a robust humoral and cellular immune response that consists in the secretion of an army of antimicrobial peptides (AMPs) in the hemolymph and the activation of *Drosophila* macrophages (plasmatocytes) (Lemaitre & Hoffmann 2007). Notably, when exposed to harmful pathogens, barrier epithelia that includes gut, trachea and epidermidis (cuticle) also have the ability to prevent infection (Bergman et al. 2017). *D. melanogaster* intestine consists in a simple ciliated epithelium layer surrounded by a muscle layer (Miguel-Aliaga et al. 2018) that has the ability to develop a proper innate immune response to intestinal bacteria, including tolerating mechanisms for beneficial microbiota (Storelli et al. 2011), similarly to mammals (Schwarzer et al. 2016). The peritrophic matrix establishes a physical barrier that isolates pathogenic bacteria and their toxins from the epithelium layer (Buchon et al. 2013). Then the *Drosophila* intestinal epithelium, at all stages, has the ability to generate an antimicrobial response. On one hand, it involves the secretion of AMPs. They are produced either upon Toll pathway activation, similarly to the MyD88-toll-like receptor pathway in mammals, reacting to Gram-positive bacteria and fungi or upon Immune deficiency (IMD) pathway activation, that shares many similarities with the Tumor Necrosis Factor (TNF) cascade, reacting to Gram negative bacteria (Capo et al. 2016; X. Liu et al. 2017). In addition to these two pathways, the fly can clear pathogenic bacteria by activating the production of microbicidal reactive oxygen species (ROS) via the Duox pathway (S.-H. Kim & W.-J. Lee 2014). Several studies showed that the *Drosophila* model recapitulates many aspects of the human intestinal pathologies (Apidianakis & Rahme 2011) and already allowed to evaluate with success the harmfulness of human pathogens such as *Mycobacterium tuberculosis* (Dionne et al. 2003), *Listeria monocytogenes* (Mansfield et al. 2003), *Vibrio cholerae* (Blow et al. 2005) or *Yersinia pestis* (Ludlow et al. 2019).

The lack of satisfactory *in vivo* model to study *S. aureus* virulence prompted us to develop an alternative *D. melanogaster* model that mimics mammalian immune responses to bacterial infections. To date, several *S. aureus* infection models have been assessed on adult flies, through systemic (via pricking in the thorax) or oral infections, but with limited -or no-infection cost on the host (Needham et al. 2004; Hori et al. 2018; Herbert et al. 2010; Ben-Ami et al. 2013; Thomsen et al. 2016; Wu et al. 2012). More specifically, among already published researches presenting *Drosophila* intestinal infection models, none of them utilized the epidemic methicillin-resistant strain *S. aureus* USA300. Thus far, *S. aureus* USA300 virulence had only been assessed by septic injury in flies, leading to animal death in a more severe way than with poorly virulent strains, *i.e. S. aureus* NCTC8325 RN1 and CMRSA6 or the colonization strain M92 (Herbert et al. 2010; Ben-Ami et al. 2013; Thomsen et al. 2016; Wu et al. 2012).

In this work, we took advantage of the *Drosophila* larval stage, when animals feed continuously and massively, to set up a new infection model based on *S. aureus* USA300 virulence. This model allowed us to characterize and follow *in vivo* at the whole organism level, bacterial fate as well as the host innate immune response triggered upon infection. We show that, through the catalase-detoxifying enzyme, *S. aureus* neutralizes intestinal epithelial ROS, hence attenuating the host immune response characterized here by the Toll pathway activation. Eventually, ROS quenching by the catalase promotes colonization of the neutral pH area of the larvae intestine and subsequently leads to animal death. We also demonstrate the colonization capacities of *Salmonella* Typhimurium *and Shigella flexneri*, suggesting that drosophila larvae could serve as a general model for the study of multiple human pathogens.

## Results

We established a 24 h infection course (**Figure 1A**) following a 30 min period where mid-L3 larvae are fed with a mix of crushed banana and bacteria (see Methods section). We observed that, after 24 h of infection, 93% of the larvae were killed by using a bacteria-enriched medium containing 2.5×10^9^ bacteria whereas lower doses (1.25×10^9^, 6.25×10^8^ or 2.5×10^8^ bacteria) killed only 62, 51 and 20% respectively of the larvae (**Figure 1B**). We next followed larval killing kinetics using wild-type *S. aureus* USA300 (WT), in comparison to the Gram-positive opportunistic entomopathogen *Micrococcus luteus* that is known to be non-pathogenic for *D. melanogaster* (Rutschmann et al. 2002). Larvae were infected with 2.5×10^9^ bacteria-enriched medium for 30 minutes and killing was followed over a 24-h period. In these conditions, *S. aureus* USA300 WT was able to kill larvae, with a drop of animal survival occurring between 12 h and 18 h **(Figure 1C)**. In contrast, *M. luteus* infection did not affect animal survival. We raised the hypothesis that animal death could be linked to bacterial load in the intestine. To avoid quantifying intestinal microbiota, we generated a *S. aureus* USA300 WT strain carrying the pRN11 plasmid that expresses a Chloramphenicol (CmR) resistance gene (de Jong et al. 2017). Comforting the survival data, a 10-fold lower bacterial number (Colony Forming Units - CFUs) was recorded in larval guts after an initial infectious dose of 2.5×10^8^ bacteria compared to 2.5×10^9^ bacteria, after 6 h, and 20-fold lower after 24 h (**Figure 1D**). We have then confirmed the absence of effective tracheal colonization. As shown in **Figure S1A**, bacterial counts remained low in the tracheal system throughout the experiment, reaching the highest CFUs count at 6 h with an average of 447 CFUs for a 2.5×10^9^ bacteria-enriched medium. Furthermore, we have shown that bacteria were not able to diffuse in the systemic compartment. As shown in **Figure S1B**, *S. aureus* USA300 WT was almost undetectable in the hemolymph as it reaches only 8 and 6 CFUs for 10 larvae respectively at 6 h and 18 h. Similar values were obtained with the non-pathogenic strain *M. luteus* (**Figure S1B**). Altogether, these data indicate that *S. aureus* USA300 WT, in this model of oral infection, is pathogenic to *D. melanogaster* larva in a dose-dependent manner, and that infection is constrained in the gut where it persists for at least 24 h.

**Figure 1.**
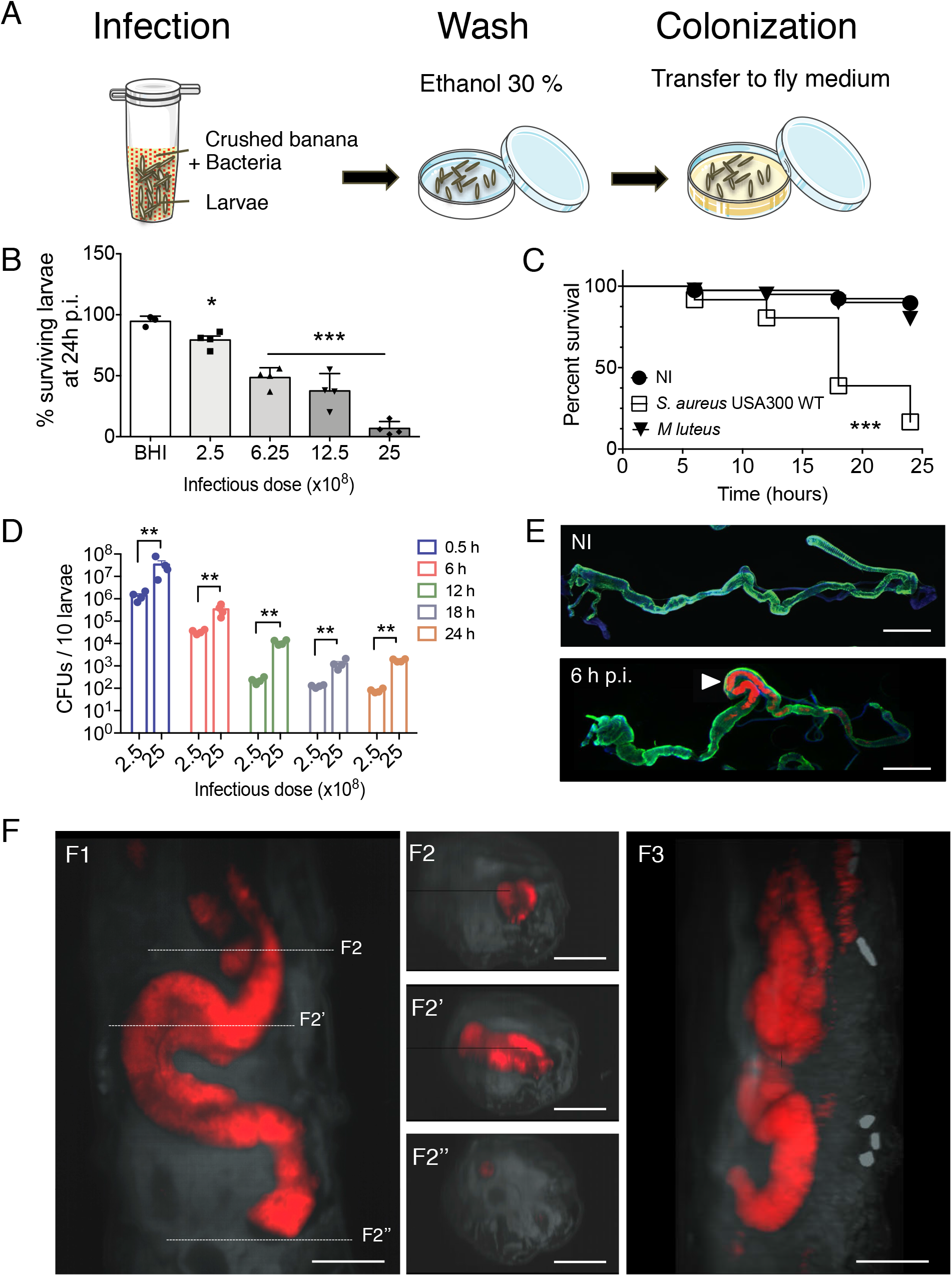
*D. melanogaster* larva is a model to study *S. aureus* USA300 virulence. (**A**) Mid-L3 larvae are placed in a microfuge tube with 100 μl of crushed banana and 100 μl of bacteria for 30 min. Then animals are briefly washed with 30% ethanol and transfer to a petri dish with fresh fly medium until further processing. **(B)** Survival of *w^1118^ D. melanogaster* larvae following 30 min oral infection with wild-type *S aureus* USA300 at the indicated infectious doses. Animals were checked 24 h after infection. Mean ± SEM, n=3 with 20 animals / point. \**P* < 0.05, \*\*\**P* < 0.001 using one-way ANOVA. **(C)** Survival of *w^1118^ D. melanogaster* larvae upon 30 min oral infection with 2.5×10^9^ of *S aureus* USA300 WT and the non-pathogenic entomopathogen *Micrococcus luteus*. Animals were followed up at 0, 6, 12, 18 and 24 h after infection. 70 animals from 3 independent experiments, \*\*\**P* < 0.001 using Kaplan-Meier test. **(D)** *w^1118^ D. melanogaster* larvae were orally infected for 30 min with 2.5×10^8^ and 2.5×10^9^ of chloramphenicol-resistant *S aureus* USA300 WT (carrying pRN11 plasmid). Bacterial counts (CFUs) in the gut were determined at 0.5, 6, 12, 18 and 24 h p.i.. Tissues were homogenized in DPBS, serially diluted and plated on BHI agar supplemented with chloramphenicol (10 μg.ml^−1^). Mean ± SEM, n=3. \*\**P* < 0.01 using two-way ANOVA. **(E)** Representative images of guts from non-infected (NI) and 6 h-infected (6 h p.i.) larvae with mCherry-*S aureus* USA300 WT (carrying pRN11 plasmid). Animals were dissected, stained with Alexa Fluor™ 488 phalloidin (green) and DAPI (blue). Scale bar = 0.5 mm. (n=2, 10 guts/experiment, for each condition) (**F)** Representative Light sheet microscope images (20X/NA0.1 objective) from posterior part (ventral view) of a larva infected with mCherry-*S. aureus* USA300 WT, at 6 h p.i. F1, F2 and F3 correspond respectively to a frontal plane (ventral view), transversal planes (reflecting arrows disposition in F1) and sagittal plane (extended view). Scale bar = 100 μm. (Experiment performed on 5 animals on 5 different inclusions).

Since oral infection of adult flies with different *S. aureus* strains, including *S. aureus* USA300, does not interfere with animal survival (Needham et al. 2004; Shiratsuchi et al. 2012; Hori et al. 2018), we hypothesized that larval death was linked to a higher number of ingested bacteria, due to their hyperphagic behavior. To confirm this, adult flies were first starved for 2 h and then placed on filters soaked with 2.5×10^9^ bacteria-enriched medium for 1 h. Neither *S. aureus* USA300 WT nor *M. luteus* oral infection affected adult flies survival (**Figure S2A**). Indeed, after 1 h feeding, bacterial counts recorded was only *ca.* 2% that recorded with larvae (*i.e.* 6.9×10^5^ bacteria/10 adult flies compared to 3.4×10^7^ bacteria/10 larvae; **Figure S2B**), suggesting that animal killing, when orally infected by *S. aureus* USA300 WT, might be dependent on the bacterial load ingested. Of note, the number of bacteria counted at day 1, in adults, is consistent with the study from Hori *et al*. (Hori et al. 2018) where they retrieved 8×10^4^ bacteria per fly gut compared to an average of 1.8×10^4^ bacteria per fly in our study. Of note, at day 1, animals display bacteria in the middle midgut (**Figure S2C**).

We next analyzed *S. aureus* localization in larval gut using fluorescence microscopy (**Figure 1E**) and Light sheet 3D imaging (**Figure 1F**and **Movie S1**). For this, we used the WT *S. aureus* USA300 strain carrying pRN11 plasmid expressing *mCherry* gene (de Jong et al. 2017) (red fluorescence). Imaging from 6 h-infected larvae with *mCherry*-expressing *S. aureus* USA300 WT revealed that bacteria were clustered in the posterior midgut (**Figure 1E-F** and **Movie S1**). This specific localization in larvae could be explained by the local gut pH, as the first half of the posterior midgut is at neutral to acidic pH, in comparison to the middle midgut that corresponds to an highly acidic region (pH < 3), and the second half of the posterior midgut to an alkaline region (pH > 10) (Shanbhag & Tripathi 2009). To verify this assumption, we tested *S. aureus* USA300 susceptibility to the pHs 3, 5, 7, 9 or 11 (**Figure S3A**). We observed that *S. aureus* USA300 was highly susceptible to basic pH 11 as well as to highly acidic pH 3. In contrast, bacteria were able to survive at pH 5 and 9, for at least 2 h and were able to multiply at pH 7. This susceptibility to environmental pH may explain the specific localization of *S. aureus* USA300 WT in the neutral region of the *Drosophila* larval gut. Interestingly, site of infection was associated with an apparent inflammation (*ca.* 1.3-fold swelling of the gut in this area compared to that of non-infected larvae, **Figure S4**). Altogether, these data show that *S. aureus* USA300 successfully colonizes *D. melanogaster* larvae upon a 24 h infection and preferentially localizes at the anterior half of the posterior midgut. This prolonged infection results in tissue inflammation and correlates with animal death.

It was previously shown that adult intestinal infection triggers the production of reactive oxygen species (ROS) through the Duox enzyme to clear invading pathogens, in complement to AMPs (Ha et al. 2005). We therefore monitored *Duox* gene transcription level in gut larvae infected with bacterial-enriched medium (2.5×10^9^ bacteria). As shown in **Figure 2A**, we observed respectively 126.5-fold and 48.9-fold increases in *Duox* transcription at 1 h and 6 h post-infection (p.i.) compared to non-infected animals. To confirm this induction of intestinal ROS in this context, we used the H_2_O_2_ specific detector 2′,7′-dichlorodihydrofluorescein diacetate (H_2_DCFDA). We observed a 20 % increase in signal detection in infected intestines, 2 h p.i., compared to non-infected intestines (**Figure 2B**). This was confirmed by live imaging as shown in **Figure 2C**. Interestingly, H_2_DCFDA fluorescence (green) was often associated with bacteria (mCherry*-S. aureus* USA300, red). We also noticed strong H_2_DCFDA fluorescence in malpighian tubules (MT) when animals were infected (**Figure 2C**,white arrows). In insects, MTs play a key role in hemolymph filtering (similar to kidneys and liver in mammals) and are intimately linked to the stress status of the fly (Davies et al. 2012). Of note, it was recently shown that MTs also play an active role during oral infection by sequestering excessive ROS and oxidized lipids (Li et al. 2020). These results demonstrate that *S. aureus* USA300 oral infection rapidly triggers H_2_O_2_ production at the intestinal epithelium, through Duox activation.

**Figure 2.**
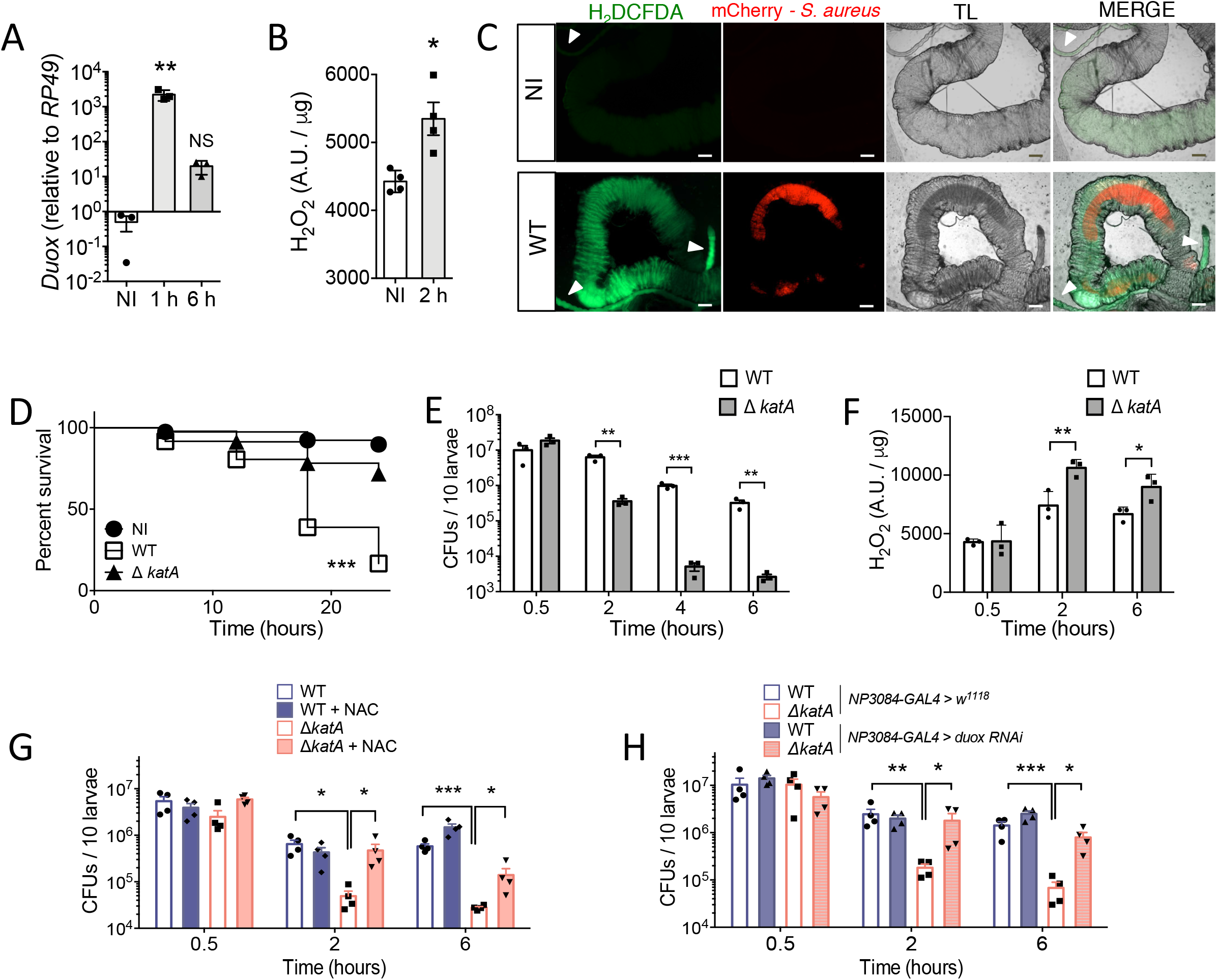
ROS quenching *in vivo* is a key mechanism for successful colonization. **(A)** *w^1118^* mid-L3 larvae were fed for 30 min with 2.5×10^9^ *S aureus* USA300 WT bacteria. Quantitative real-time PCR analysis on *Duox* transcripts was done with total RNA extracts from guts (15 animals, n=3) recovered at 1 h and 6 h p.i. Bar graph data are presented related to *RP49*. Mean ± SEM, n=3, NS= non-significant, \*\**P* < 0.01 against NI using one-way ANOVA. **(B)** *w^1118^* mid-L3 larvae were fed for 30 min with 2.5×10^9^ *S aureus* USA300 WT bacteria. Generation of intestinal H_2_O_2_ was measured with the H_2_DCFDA dye (10μM) on non-infected samples and at 2 h p.i.. Mean ± SEM, n=4, \**P* < 0.05 using Mann-Whitney test. **(C)** Representative live imaging of posterior midgut from non-infected larvae (NI) and orally infected larvae (2 h p.i., mCherry-*S. aureus* USA300 WT, red). Intestines are dissected, treated with H_2_DCFDA (10 μM, green) for 15 min and imaged with an epifluorescence microscope. TL = Transmitted light. White arrows indicate malpighian tubules. Scale bar = 10 μm. (n=3, 10 guts/experiment, for each condition) **(D)** Survival of *w^1118^ D. melanogaster* larvae following 30 min oral infection with wild-type *S aureus* USA300 WT or *S aureus* USA300 *ΔkatA* with 2.5×10^9^ bacteria. Experiment was followed up until 24 h after infection. 75 animals pooled from 3 independent experiments, \*\*\**P* < 0.001 using Kaplan-Meier test. **(E)** *w^1118^ D. melanogaster* larvae were orally infected for 30 min with 2.5×10^9^ bacteria of chloramphenicol-resistant *S aureus* USA300 WT or *S aureus* USA300 *ΔkatA* (carrying pRN11 plasmid). Bacterial counts (CFUs) were determined at 0.5, 2, 4 and 6 h p.i.. After homogenization and serial dilution, samples were plated on BHI supplemented with chloramphenicol (10 μg.ml^−1^). Mean ± SEM, n=3, \*\**P* < 0.01, \*\*\**P* < 0.001 using two-way ANOVA. (**F**) *w ^1118^* mid-L3 larvae were fed for 30 min with chloramphenicol-resistant *S. aureus* USA300 WT or *S. aureus* USA300 *ΔkatA* at the infectious dose of 2.5×10^9^ bacteria. Intestinal ROS titer was measured at 0.5, 2 and 6 h p.i. After dissection, intestines were homogenized in 400 μl DPBS and treated with H_2_DCFDA (10μM) for 30 min. Fluorescence was measured at 490nm. Mean ± SEM, n=3, \**P* < 0.05, \*\**P* < 0.01 using two-way ANOVA. **(G)** *w^1118^D. melanogaster* mid-L3 larvae were orally infected for 30 min with chloramphenicol-resistant *S. aureus* USA300 WT or *S. aureus* USA300 *ΔkatA* (carrying pRN11 plasmid) at the infectious dose of 2.5×10^9^ bacteria. Then animals were transferred to fresh fly medium supplemented, or not, with NAC (1mM). Bacterial counts (CFUs) were determined at 0.5, 2 and 6 h p.i. After homogenization and serial dilution, samples were plated on BHI supplemented with chloramphenicol (10 μg.ml^−1^). Mean ± SEM, n=4, \**P* < 0.05, \*\*\**P* < 0.001 using two-way ANOVA. **(H)** *NP3084-GAL4* > *w^1118^ and NP3084-GAL4* > *Duox RNAi* larvae were orally infected for 30 min with chloramphenicol-resistant *S. aureus* USA300 WT or *S. aureus* USA300 *ΔkatA* (carrying pRN11 plasmid) at the infectious dose of 2.5×10^9^ bacteria. Bacterial counts (CFUs) were determined at 0.5, 2 and 6 h p.i.. After homogenization and serial dilution, samples were plated on BHI supplemented with chloramphenicol (10 μg.ml^−1^). Mean ± SEM, n=4, \**P* < 0.05, \*\**P* < 0.01, \*\*\**P* < 0.001 using two-way ANOVA.

As H_2_O_2_ generation through Duox enzyme is a key mechanism to control pathogen load (K.-A. Lee et al. 2015), we focused our attention on the contribution of the catalase, encoded by the *S. aureus katA* gene, in animal infection. Catalase enzyme detoxifies hydrogen peroxide by converting it to water and oxygen molecules. *katA* gene constitutes the unique catalase encoding gene in *S. aureus* (Horsburgh et al. 2001; Beavers & Skaar 2016). First, we evaluated *D. melanogaster* survival to *S. aureus* USA300 *ΔkatA* oral challenge. For this, we used a mutant from Nebraska Transposon Mutant Library that carries a transposon insertion in the *katA* gene (NE1366 from BEI resource). We first confirmed that the *ΔkatA* mutant strain had no growth defect in liquid broth (BHI, **Figure S5A**) and was more sensitive to H_2_O_2_ than the WT strain (15 mM H_2_O_2_ in DPBS, **Figure S5B**). As shown in **Figure 2D**, we observed that *S. aureus* USA300 *ΔkatA* killed larvae to a much lesser extent than the WT strain. This difference in larval survival was correlated with a 17.5-, 190- and 122-fold decrease in intestinal *ΔkatA* mutant CFUs compared to WT CFUs, respectively at 2, 4 and 6 h p.i. (**Figure 2E**). Supporting the notion that this higher bacterial clearance could be related to a defect in quenching H_2_O_2_, we observed a significant increase in ROS amount, by H_2_DCFDA measurement, in larval intestines infected with the *ΔkatA* strain in comparison to animals infected with the WT strain, with a respective increase of 43% and 34% of fluorescence intensity at 2 and 6 h p.i. (**Figure 2F**). This result suggests that *S. aureus* USA300 *ΔkatA* is defective for H_2_O_2_ quenching. Then, we confirmed that bacterial persistence into larval intestine and bacterial ability to kill larvae are closely related to ROS content. For this, we evaluated bacterial CFUs of WT and *ΔkatA* strains in flies fed with N-acetyl-L-cysteine (NAC, 1mM), an antioxidant drug that was shown to quench H_2_O_2_ molecules (Aruoma et al. 1989). We observed that NAC counteracted the deleterious intestinal environment for the *ΔkatA* strain as NAC abolished *ΔkatA* defect compared to WT, at 2 h p.i., and promoted a 5-fold increase in CFUs count of the *ΔkatA* 6 h p.i. (**Figure 2G**). In parallel, we have tested WT and *ΔkatA* strains survival in *NP3084-GAL4 > Duox-RNAi* larvae, that are defective for *Duox* expression specifically in the intestine. *NP3084-GAL4* (or *MyoD1-GAL4*) drives predominantly gene expression in larval midgut, in enterocytes (Nehme et al. 2007). Notably, larvae whose *Duox* expression was abolished in the midgut (*NP3084-GAL4 > Duox-RNAi*) showed a significant 9.8- and 11.7-fold increases in CFUs counts for the *ΔkatA* strain, respectively at 2 and 6 h p.i., in *NP3084-GAL4 > Duox-RNAi* compared to *NP3084-GAL4 > w^1118^* larvae. In contrast, *S. aureus* USA300 WT strain showed non-significant 0.8 and 1.7-fold changes in CFUs counts in *NP308-GAL4 > Duox-RNAi* compared to *NP3084-GAL4 > w^1118^* larvae (**Figure 2H**). Together, these results indicate that oral infection induces H_2_O_2_ generation from epithelial barrier that acts as a key mechanism to control the growth of pathogen. Besides, the catalase activity is paramount to *S. aureus* resistance to host response through H_2_O_2_ quenching.

As other Gram-positive bacteria, *S. aureus* is known to induce the Toll pathway, one key innate immune signaling in *D. melanogaster*, through its lysine-type peptidoglycan (Buchon et al. 2014). This prompted us to test the expression of the *Drosomycin* gene (*Drs,* which encodes an antimicrobial peptide and which is one of the main read-outs of the Toll pathway in *D. melanogaster*) in *yw* wild type larvae and the derivative *spz^rm7^* mutated line (larvae lacking the expression of the Toll ligand spätzle) when infected with the *S. aureus* USA300 WT strain. As shown in **Figure 3A**, we observed a significant 126-fold increase in intestinal *Drs* expression, in comparison to non-infected conditions (using a 2.5×10^9^ bacteria-enriched medium), in *yw* flies. This activation was proportional to the initial bacterial load as a 10-fold lower infectious dose (2.5×10^8^) induced only a 16-fold increase in *Drs* gene transcription. Notably, using *spz^rm7^* larvae considerably reduced *Drs* transcripts amount, even using 2.5×10^9^-enriched medium, suggesting that *Drs* activation is almost exclusively controlled by the Toll pathway. In *Drosophila,* links between ROS and the Toll/NF-κB pathway has already been established. Under wasp infestation (at larval stage), the lymph gland (the main hematopoietic organ) undergoes a burst of ROS in the posterior signaling center, resulting in Toll pathway activation and whose purpose is to redirect hemocyte progenitors differentiation into lamellocytes subtype (Louradour et al. 2017). This led us to wonder if H_2_O_2_ generated during the infection could play a role in Toll pathway activation in the intestine. For this, we have first tested the direct effect of H_2_O_2_ on intestinal *Drs* expression, a reporter gene for Toll pathway activation. Interestingly, animals treated with H_2_O_2_ (0.5 % in fly medium) for 2 h showed a 15-fold induction of the *Drs* expression in the gut (**Figure 3B**). We then evaluated the expression of *Drs* gene in flies infected with the WT and *ΔkatA* strains, fed with NAC or not. We first observed, at 6 h p.i, that the WT and the *ΔkatA* strains respectively induced a 91- and 169-fold increase in *Drs* expression relative to non-infected condition. Notably, at 6 h p.i., our results (**Figure 2B**) showed a 122-fold decrease in *S. aureus* USA300 *ΔkatA* strain numeration compared to WT strain, suggesting that other factors than the bacteria themselves modulate *Drs* expression. Feeding animals with NAC induced relative 1.8 and 3.5-fold decreases in *Drs* expression in larval intestine, 6 h p.i. when infected respectively with the *ΔkatA* or the WT strains (**Figure 3C**). These results highlight the close relationship between ROS generation and immune response activation in host when infected by *S. aureus*. Indeed, ROS neutralization by bacterial catalase enzyme counteracts the dual deleterious effects of ROS molecules either through direct damages, or through Toll pathway activation with Drosomycin AMP secretion.

**Figure 3.**
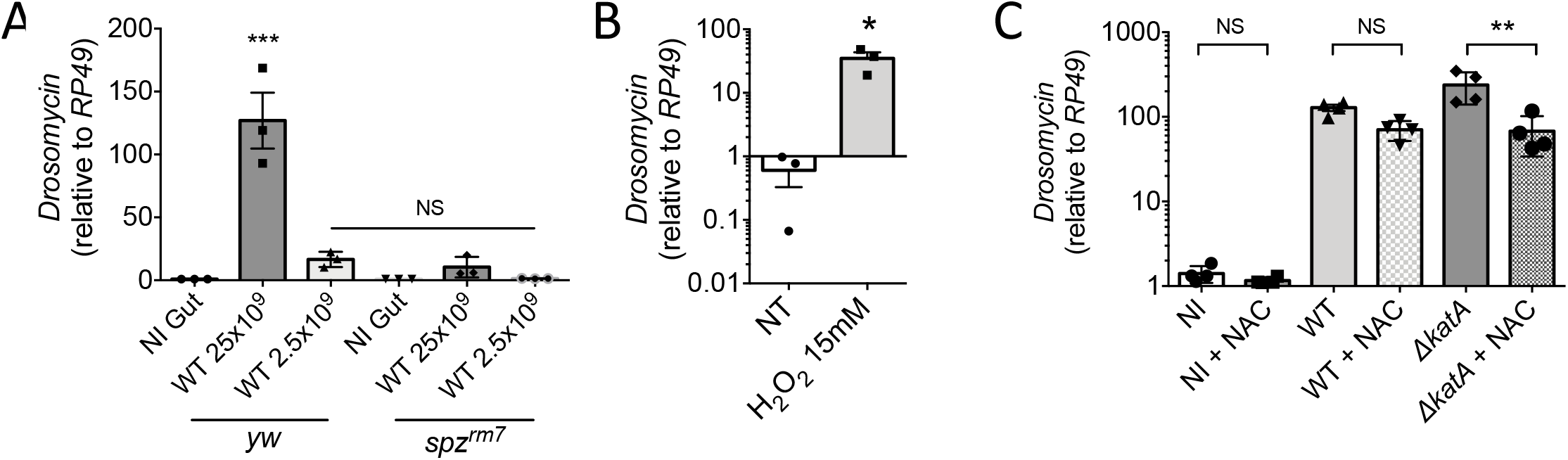
S. aureus USA300 modulates antimicrobial response by neutralizing intestinal ROS. **(A)** *yw* and *yw;;spz^rm7^* mid-L3 larvae were fed for 30 min with bacteria at the infectious doses of 2.5×10^8^ and 2.5×10^9^ bacteria. Quantitative real-time PCR analysis on *Drosomycin* transcripts was done with total RNA extracts from guts recovered at 6 h p.i.. Bar graph data are presented related to *RP49*. Mean ± SEM, n=3, NS = non-significant, \*\*\**P* < 0.001 using one-way ANOVA. **(B)** *w^1118^* mid-L3 larvae were fed for 2 h with fly medium supplemented with stabilized H_2_O_2_ (0.5%). Guts were dissected for quantitative real-time PCR analysis on *Drosomycin* transcripts. Transcripts levels were normalized to the corresponding *RP49* levels. Mean ± SEM, n=3, \**P* < 0.05 using Mann-Whitney test. **(C)** *w^1118^* mid-L3 larvae were orally infected for 30 min with *S aureus* USA300 WT or *S aureus* USA300 *ΔkatA* at the infectious dose of 2.5×10^9^ bacteria. Then animals were transferred to fresh fly medium supplemented, or not, with NAC (1mM). At 6 h p.i., guts were dissected for quantitative real-time PCR analysis on *Drosomycin* transcripts. Data were normalized to the corresponding *RP49* levels. Results were compared to non-infected larvae transferred on supplemented NAC medium (NI+NAC) or not (NI). Mean ± SEM, n=4, NS = not-significant, \*\**P* < 0.01 using one-way ANOVA.

We have shown above that the flies larvae represent an appropriate and suitable model to study host-pathogen interaction with the multiresistant strain *S. aureus* USA300. To evaluate the larval model polyvalence and confirm its ease of implementation, we have tested larvae infection with two human enteric pathogens: *Salmonella enterica* serovar Typhimurium and *Shigella flexneri* (see Materials and Methods section for details). Notably we observe significant larval death when fed with 2.5×10^9^ bacteria, in similar condition than *S. aureus* infection (see Material and Methods section). After 24 h of infection, 53.8 % and 46.6 % of the larvae are killed respectively when fed with *S.* Typhimurium and *S. flexneri* enriched medium (**Figure 4A**). After 30 min feeding, larvae are infected with 8.2 × 10^6^ and 7.9 × 10^6^ bacteria per 10 animals, respectively with *S.* Typhimurium and *S. flexneri*. At 6 h post-infection, counts reach respectively 5 × 10^5^ and 2 × 10^5^ bacteria (**Figure 4B**). Using this model we have confirmed that these two pathogens trigger *Drosophila* immune response as we have observed a significant production of intestinal H_2_O_2_ at 2 h when infected with *S.* Typhimurium (43 % increase) and *S. flexneri* (57 % increase) (**Figure 4C**). This was correlated with a significant increase in the expression of the antimicrobial peptide *Diptericin* (*Dpt*) gene, that is dependent on the Gram-negative sensitive Immune deficiency pathway (Lemaitre & Hoffmann 2007). We observed respectively 42.9 and 37.9-fold increases in *Dpt* expression at 6 h post-infection (**Figure 4D**). Interestingly, by using DsRed expressing strains, we observed that, at 6 h post-infection, each strain localized preferentially at the anterior and middle midgut (white arrows) where pH values vary from neutral to acidic values (Shanbhag & Tripathi 2009). A trait that can be explained by their ability to survive acidic environment (Lin et al. 1995). We confirmed *S.* Typhimurium and *S. flexneri* lower susceptibility to pH 5 after 2 h of treatment, in comparison to *S. aureus* (Figure S6). After 2 h in BHI broth adjusted at pH 5, the number of *S. aureus* CFUs was 25-fold lower than that recorded after pH 7 treatment. In contrast, for *S.* Typhimurium and *S. flexneri*, the values recorded at pH5 were only 5-to 3-fold lower, respectively than those at pH7.

**Figure 4.**
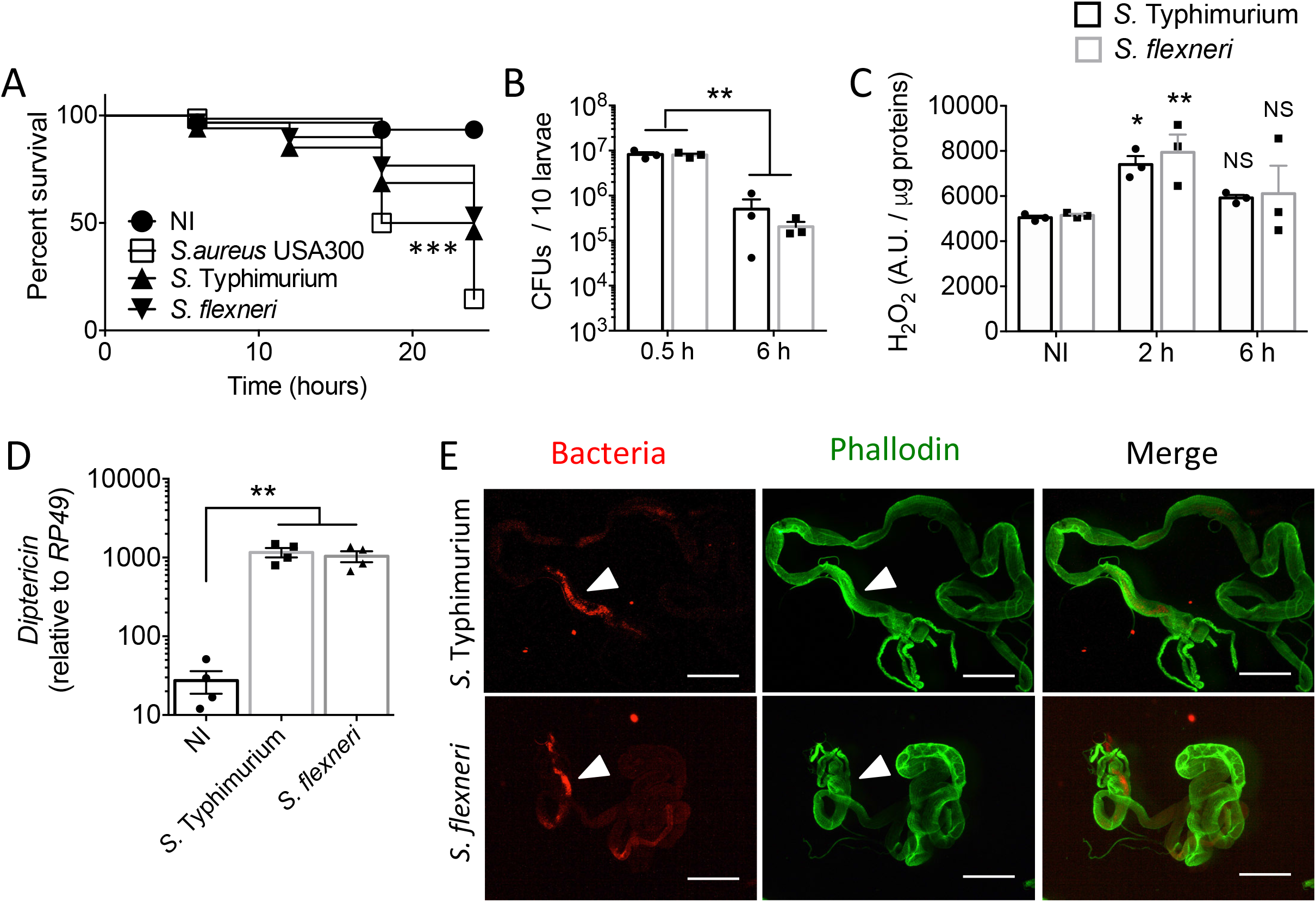
D. melanogaster larva can serve as a model host for human enteric pathogens. **(A)** Survival of *w^1118^ D. melanogaster* larvae upon 30 min oral infection with 2.5×10^9^ of *S. aureus* USA300, *S.* Typhimurium or *S. flexneri*. Animals were followed up at 0, 6, 12, 18 and 24 h after infection. 65 to 70 animals from 3 independent experiments were pooled, \*\*\**P* < 0.001 using Kaplan-Meier test. **(B)** *w^1118^ D. melanogaster* larvae were orally infected for 30 min with 2.5×10^9^ of chloramphenicol-resistant *S aureus* USA300 (carrying pRN11 plasmid) or ampicillin-resistant *S.* Typhimurium and *S. flexneri.* Bacterial counts (CFUs) in the gut were determined at 0.5 and 6 h p.i.. Tissues were homogenized in DPBS, serially diluted and plated on BHI agar supplemented with chloramphenicol (10 μg.ml^−1^) or ampicillin (100 μg.ml^−1^). Mean ± SEM, n=3. \*\**P* < 0.01 using two-way ANOVA. **(C)** *w^1118^* mid-L3 larvae were fed for 30 min with 2.5×10^9^ *S.* Typhimurium or *S. flexneri* bacteria. Generation of intestinal H_2_O_2_ was measured with the H_2_DCFDA dye (10μM) on non-infected samples, at 2 and 6 h p.i.. Mean ± SEM, n=3, \**P* < 0.05 and \*\**P* < 0.01 using Mann-Whitney test. **(D)** *w^1118^* mid-L3 larvae were fed for 30 min with bacteria at the infectious doses of 2.5×10^9^ bacteria. Quantitative real-time PCR analysis on *Diptericin* transcripts was done with total RNA extracts from guts recovered at 6 h p.i.. Bar graph data are presented related to *RP49*. Mean ± SEM, n=4, \*\**P* < 0.01 using one-way ANOVA. **(E)** Representative images of guts from 6 h-infected larvae with DsRed *S.* Typhimurium or *S. flexneri*. Animals were dissected, stained with Alexa Fluor™ 488 phalloidin (green) and DAPI (blue). Scale bar = 0.5 mm. (n=3, 25 guts in total for each condition)

## Discussion

We present here an *in vivo Drosophila* larvae model that allows to easily and rapidly follow simultaneously bacterial infection and host innate immune responses with three human pathogens: *S. aureus*, *S.* Typhimurium and *S. flexneri.* Mammalian models, and specifically mice, are predominant in the study and identification of *S. aureus* key virulence factors (H. K. Kim et al. 2014). The use of invertebrates model has also shown great potential for dissecting complex host-pathogen interactions (Kurz & Ewbank 2000; García-Lara et al. 2005; Edwards & Kjellerup 2012). Among others, *Drosophila melanogaster* has many advantages as an experimental system, displaying remarkable high innate immunity homology with mammals in addition to the available genetic tools and breeding facilities.

We observed that upon oral *Drosophila* larvae infection with *S. aureus,* bacteria reach and establish in the first half of the posterior part of the larval intestine, possibly due to the neutral pH specifically encountered at this site. This localized colonization was associated with intestine enlargement, as observed earlier with the invertebrate model *Caenorhabditis elegans* (Irazoqui et al. 2010). Notably, we also observed that infection with 2.5×10^9^ bacteria finally kills larvae in a significant manner after 24 h. In contrast to a previous study performed with non antibiotics-resistant *S. aureus* strains and that did not identify a killing effect from bacteria at the adult stage (Hori et al. 2018), here we suggest that the killing phenotype observed in larvae is due primarily to the amount of ingested bacteria. In addition, the epidemic strain USA300 carries an hypervirulent phenotype characterized by the expression of multiple toxins (such as the enterotoxins K and Q, and the Panton-Valentine Leukocidin pore forming toxin) and the arginine catabolic mobile element (ACME) that displays adhesive properties and improves bacterial colonization (Thurlow et al. 2012; Jones et al. 2014). All these specificities could play an important role for successful intestinal establishment.

It was recently shown that larvae orally infected with *Erwinia carotovora carotovora 15* (*Ecc15*), *Pseudomonas aeruginosa* or *Pseudomonas entomophila* are more susceptible to pathogens in comparison to adult flies infected with similar doses (Houtz et al. 2019). Adult intestine undergoes basal turnover characterized by intestinal stem cells (ISCs) proliferation that differentiate into intermediate progenitor cells named enteroblasts (EBs), then into enterocytes (ECs) or enteroendocrine cells (EEs) (Hung et al. 2020). Upon infection with the Gram-negative pathogens *Ecc15* (Buchon et al. 2009; Buchon et al. 2010) or *P. entomophila* (Jiang et al. 2009), compensatory mechanisms respectively activated by the Epidermal Growth Factor Receptor (EGFR) and the JAK/STAT pathways, initiate a strong mitotic response in the midgut, without modifying ISCs number. This phenomenon is complementary to the intestinal antimicrobial response and essential to resist infection. Interestingly, it was also shown that the IMD pathway plays a key role in ECs shedding during infection, also favoring epithelial turnover (Zhai et al. 2018). In contrast, *Drosophila* larvae are devoted of ISCs and, upon *Ecc15* intestinal infection, rely on adult midgut progenitors found in islets surrounded by peripheral cells (Houtz et al. 2019). These progenitor cells differentiate into ECs however the authors raise the hypothesis that these cells are insufficient in number to meet the need of both intestinal repair and antimicrobial response.

Interestingly, we also found that *S. aureus* infection triggers ROS production in the intestine, early during infection course and in a transient way. The *Drosophila* genome encodes one Duox enzyme whereas two Duoxs homologs are identified in mammals (Ewald 2018). In flies, it was shown that Duox enzyme can be activated by pathogens-derived uracil, unlike commensal bacteria that do not secrete this molecule (Valanne & Rämet 2013). Upon intestinal infection, released uracil induces the Hedgehog signaling pathway to maintain high Cadherin 99C level, leading to a PLCβ/PKC/Ca2+ dependent Duox activation (K.-A. Lee et al. 2015). Enteric infection is also responsible for a metabolic shift associated with a pronounced lipid catabolism in enterocytes, subsequently leading to cellular NADPH increase and Duox activation (K.-A. Lee et al. 2018). Of note, *S. aureus* USA300 is capable of generating uracil through pyrimidine metabolism (Kyoto Encyclopedia of Genes and Genomes pathway). We herein showed that an increase of *Duox* gene transcription, associated with an increase of ROS generation (measured by the dichlorofluorescin diacetate specific probe), occurred in the first hours of infection. These data somewhat contradict an earlier work by Hori *et al.* (Hori et al. 2018), reporting that *S. aureus* Drosophila feeding did not induce ROS production. This apparent discrepancy could likely to be due to the ROS quantitation method used in both work. Hori and colleagues used hydro Cy3 to quantify ROS amount, a compound for which measurement may be influenced by mitochondrial membrane potential (Zhdanov et al. 2017) which are modified during cell bacterial infection (Tiku et al. 2020). This discrepancy could also be due to the method used to dissect larval intestine. In insects, malpighian tubules play a key role in detoxification and hemolymph filtering (similar to kidneys and liver in mammals). They are intimately linked to the stress status of the fly (Davies et al. 2012) and it was recently shown that malpighian tubules play an active role during oral infection by sequestering excessive ROS and oxidized lipids (Li et al. 2020). Including them during dissection could greatly affect final results by hiding the specific intestinal ROS signal. In another model of orally infected black soldier flies, *S. aureus* was shown to be able to induce *Duox* gene expression as well as increasing H_2_O_2_ concentration, also in short time manner (Yu et al. 2019). Overall, this work confirms the importance of generating intestinal oxidative stress to clear colonizing pathogens as well as the necessity for the bacterium to acquire efficient oxidative stress resistant systems. Our results demonstrate that *S. aureus* USA300 *catalase* gene is necessary to increase bacterial virulence *in vivo* and assess its colonization capacities. Of note, *S. aureus* catalase gene importance has previously been shown *in vitro* during intracellular infection in murine macrophages or *in vivo* through intraperitoneal injection with a clinical bovine strain (Martínez-Pulgarín et al. 2009), where bacteria were directly injected.

This work highlighted the link between ROS production and Toll signaling activation in the gut following *S. aureus* USA300 exposure. One may assume that this mechanism could potentiate the host immune response against harmful pathogens such as *S. aureus.* In addition to already established link between ROS and Toll pathway triggering in *D. melanogaster,* it was reported that *Wolbachia* infected mosquitoes had an increase in *Duox2* transcription and this was sufficient to induce transcription of the Toll pathway susceptible AMPs cecropins and defensins (Pan et al. 2012). In *Drosophila,* ROS and the Toll/NF-κB are also closely linked. Under wasp infestation (at larval stage), the lymph gland (the main hematopoietic organ) undergoes a ROS burst in the posterior signalling center, resulting in Toll activation and whose purpose is to redirect hemocyte progenitors differention into lamellocytes subtype (Louradour et al. 2017). Researches in mammals suggest that ROS can alter iκB kinase complex (IKK) activity in the cytoplasm or NF-κB DNA binding capacity in the nucleus (Morgan & Z.-G. Liu 2011). These observations highlight the need for the bacteria to consistently control the host clearance strategy, by simultaneously acting on the immune response and the ROS pool.

Overall, our observations suggest that the drosophila larval infection model could serve as an easy-to-manipulate system to study host innate immune responses triggered upon infection with human bacterial pathogens. They support the potential of invertebrate models as promising alternatives to mammalian models.

## Materials and Methods

### Bacterial strains

The epidemic clone *S. aureus* USA300-LAC (designated as *S. aureus* USA300 WT) as well as its isogenic derivative *S. aureus* USA300-Δ*katA,* were provided by the Biodefense and Emerging Infections Research Resources (BEI). Importantly, *S. aureus* USA300 is the leading epidemic lineage, among antibiotics resistant and sensitive *S. aureus* strains, found in United States and Europe. They were grown in Brain Heart Infusion (BHI) broth, at 37°C. Chloramphenicol resistant (CmR) and mCherry fluorescent strains were generated by introducing the pRN11 plasmid (de Jong et al. 2017) by electroporation, with the following settings: 2,450 V, 100 Ω, 25 μF, time constant = 2.3–2.5 ms. *S. aureus* growth was performed in Brain Heart Infusion (BHI) broth at 37°C and *Micrococcus luteus* was grown in Luria-Bertani (LB) broth at 30°C. *Salmonella* Typhimurium SL1344 was grown in LB broth or agar, at 37°C, and *Shigella flexneri* was grown in Tryptic Soy Broth or Agar (TSA) supplemented with Congo Red dye (final concentration 0.01%) to induce type 3 secretion system (T3SS) dependent secretion of virulence factors (Parsot et al. 1995). Only pigmented colonies from TSA plates were used to prepare liquid cultures.

All strains are defined in the **Table 1**.

**Table 1.**

| Strains or plasmid | Relevant characteristics | Source or reference |
| --- | --- | --- |
| Strains |  |  |
| *Staphylococcus aureus* USA300 WT | MRSA, JE2 isolate (ST8), Ery<sup>R</sup> | BEI resources |
| *Staphylococcus aureus* USA300 WT – pRN11 (mCherry) | MRSA, JE2 isolate (ST8), Ery<sup>R</sup>, Cm<sup>R</sup>. mCherry expressing. | This work |
| *Staphylococcus aureus* USA300 $\Delta katA$ | MRSA, JE2 isolate (ST8), Ery<sup>R</sup> Transposon insertion in *catalase* gene, position 1350703, reverse orientation. | BEI resources (NE1366) |
| *Staphylococcus aureus* USA300 $\Delta katA$ – pRN11 mCherry | MRSA, JE2 isolate (ST8), Ery<sup>R</sup>, Cm<sup>R</sup> Transposon insertion in *catalase* gene, position 1350703, reverse orientation. mCherry expressing. | This work |
| *Micrococcus luteus* |  | Gift from D. Ferrandon laboratory |
| *Salmonella* Typhimurium – pGG2 DsRed | *Salmonella enterica* serovar Thyphimurium SL1344, transformed with pGG2 expressing DsRed under the rpsM promoter, Amp<sup>R</sup>. | (Voznica et al. 2018) |
| *Shigella flexneri* -pMW211 pDsRed | *Shigella flexneri* strain M90T Sm (serotype 5a), DsRed expressing, Amp<sup>R</sup> | (Sansonetti et al. 1982; Sørensen et al. 2003) |
| Plasmid |  |  |
| pRN11 | sarA p1-mCherry, vector backbone pCM29 in *E. coli* DC10b, Amp<sup>R</sup> in *E. coli* and Cm<sup>R</sup> in Gram positive bacteria. | (de Jong et al. 2017) |

### Drosophila stocks and rearing

*Drosophila melanogaster* was maintained on a fresh medium prepared with the Nutrifly Bloomington formulation (Genesee Scientific, San Diego, CA, USA), supplemented with 64 mM propionic acid and dried yeast. N-acetyl-L-cysteine (NAC) supplemented medium were prepared at the final concentration of 1 mM (A9165, Sigma (Shaposhnikov et al. 2018)).

All *Drosophila* stocks are defined in the **Table 2**.

**Table 2.**

| *Drosophila melanogaster* lines | Source | Identifier |
| --- | --- | --- |
| *w*<sup>1118</sup> (control line) | D. Ferrandon |  |
| *ywDD1*; (control line) | D. Ferrandon | (Ferrandon et al. 1998) |
| *yw,drs-GFP ,dipt-LacZ,;;spz<sup>rm7</sup>/TM6c*<br>(*ywDD1;+;spz<sup>rm7</sup>/TM6c*) | D. Ferrandon | (Ferrandon et al. 1998) |
| *w;NP3084-GAL4;+* | W.J. Lee | DGRC (113094) |
| *w;UAS-Duox RNAi/CYO ;* | W.J. Lee | (Ha et al. 2005) |

### Infection experiments

Oral infections were performed on mid-L3 larvae (3.5 days after egg-laying). For each test, animals were placed in a 2 mL microfuge tube filled with 100 μl of crushed banana (without yeast) and 100 μl of overnight bacterial culture, for 30 min. Bacterial infectious dose were adjusted by measuring culture turbidity at OD_600_. Animals were blocked by a foam plug to be sure they remain in the bottom of the tube for the whole infection time. After 30 min, they are washed briefly in ethanol 30% and placed in petri dish with fresh fly medium without yeast. Infections and waiting times were performed at 29°C. Larvae were dissected at indicated time points for RT-qPCR analyses, bacterial counts and ROS quantification.

For adults oral infection, 5-7 days old adults were starved for 2 h in empty vials at 25°C. After starvation, flies were flipped to an infection vial with medium, completely covered with a Whatman paper disk. The disk was soaked with 100 μl of a 5% sucrose solution supplemented or not with bacteria at the indicated infectious doses. After 30 min of oral infection, flies were flipped to fresh fly medium without yeast (changed every day).

### CFUs counts

To count living bacteria in gut larvae and trachea, and to avoid bacterial contaminant when plating, we used pRN11-carrying bacteria. pRN11 plasmid confers resistance to chloramphenicol antibiotics. At indicated time points, larvae were dissected (at least 10 animals per point) and guts homogenized in 400 μl of Dulbecco’s phosphate-buffered saline (DPBS, Gibco, ThermoFisher Scientific, MA, USA) with a Mikro-Dismembrator S (Sartorius stedim, Aubagne, France). Samples were serially diluted and plated on BHI with chloramphenicol (10 μg.ml^−1^, Sigma-Aldrich, Mi, USA). For CFUs count from hemolymph, animals were briefly washed in ethanol 70%, rinsed in sterile DPBS and bled into a 200 μl DPBS drop on slide. Samples were directly plated on BHI agar plates for *S. aureus* counts or LB agar for *M. luteus.*

### Bacterial survival assay

BHI pH was adjusted with sodium-chloride or hydrochloric acid solutions at the selected conditions: pHs 3, 5, 7, 9 and 11. Fresh bacterial cultures that reached OD_600_ of 0.3-0.6 were washed one time with PBS and then diluted in the different buffers to reach the concentration of 2×10^7^ bacteria/mL. At the indicated time points, 50 μl from each culture was sampled, serially diluted and plated on BHI agar.

### ROS quantification and visualization

#### ROS quantification

Amount of ROS in dissected guts (from 10 animals) was estimated using 2’,7’-Dichlorodihydrofluorescein diacetate (H_2_DCFDA, C6827, ThermoFisher Scientific, MA, USA), following manufacturer’s instructions. For larval guts dissection, we carefully removed malpighian tubules as they can strongly influence ROS level, then tissues were homogenized in H_2_DCFDA mix. Fluorescence was measured 30 min after mix preparation in multiplate reader Berthold TriStar LB941 (Berthold France SAS, Thoiry, France). Results were normalized to total protein for each sample. Proteins concentration was quantified using Pierce BCA colorimetric assay (Life Technologies, Ca, USA), following manufacturer’s instructions.

#### ROS visualization

Guts were dissected at indicated time on glass slides, incubated in H_2_DCFDA (10 μM) for 15 min and live-imaged with a Zeiss Axioimager Z2 Apotome microscope.

### Larval imaging

#### Whole gut stainings

Guts were dissected in PBS, fixed for at least 1 h at room temperature in 4% paraformaldehyde in PBS and permeabilized in PBS+0.1% Triton X-100 for 30 min. They were stained with Bodipy™ 493/503 at the dilution 1/100 (D3922, ThermoFisher Scientific, MA, USA) for 1 h, stained with DAPI at the dilution 1.43 μM for 10 min, washed with PBS and mounted in Mowiol 4-88 (17951-500, Biovalley, France).

#### Light Sheet Fluorescence Microscopy (LSFM)

For sample preparation, animals were firstly fixed in ScaleCUBIC-1 (reagent-1) for at least 4 days and cleared in ScaleCUBIC-2 (reagent-2) for at least 2 days according to Susaki *et al.* (Susaki et al. 2015). Briefly, to prepare 500g of reagent-1 solution, 125g of urea and 156g of 80 wt% Quadrol are dissolved in 144g of dH_2_0. After complete dissolution under agitation, we add 75g of Triton-X100 and then degas the reagent with vacuum desiccator (~0.1 MPa, ~30 min) (Susaki et al. 2015). Then samples were cleared with ScaleCUBIC-2 (reagent-2). To perform the LSFM imaging, samples were embedded in 4% low-melting agarose (Thermo Fisher Scientific, France) dissolved in R2 medium, by using a glass cylindrical capillary, and allow embedding overnight. Images were acquired with a Lightsheet Z.1 Microscope (Carl Zeiss, Germany) equipped with a Plan-Apochromat 20x/NA1 R2-immersion objective lens with left and right illumination.

### Quantitative Reverse Transcription PCR

For mRNA quantification, dissected guts (from 15 animals) were collected at indicated time points and homogenized with a Mikro-Dismembrator S (Sartorius stedim, Aubagne, France). Total RNA was isolated using TRIzol reagent and dissolved in RNase-free water. Five hundred nanograms total RNA was then reverse-transcribed in 20 μl reaction volume using the LunaScript RT SuperMix Kit (E3010, New England Biolabs, MA, USA). Quantitative PCR was performed by transferring 2 μl of the RT mix to the qPCR mix prepared with Luna Universal qPCR Master Mix (M3003, New England Biolabs, MA, USA), according to the manufacturer’s recommendations. All the primers used for this experiment are defined in the **table 3** and their amplification efficiency was checked before any further analysis. Reactions were performed on a 7900HT Fast Real-Time PCR System (Applied Biosystems) according to the standard settings of the system software. The thermal cycling conditions were: initial denaturation at 95°C for 1 min followed by 40 cycles of 95°C for 15 s and 60°C for 30 s. We used relative quantification with normalization against the reference gene *RP49.*

**Table 3.**
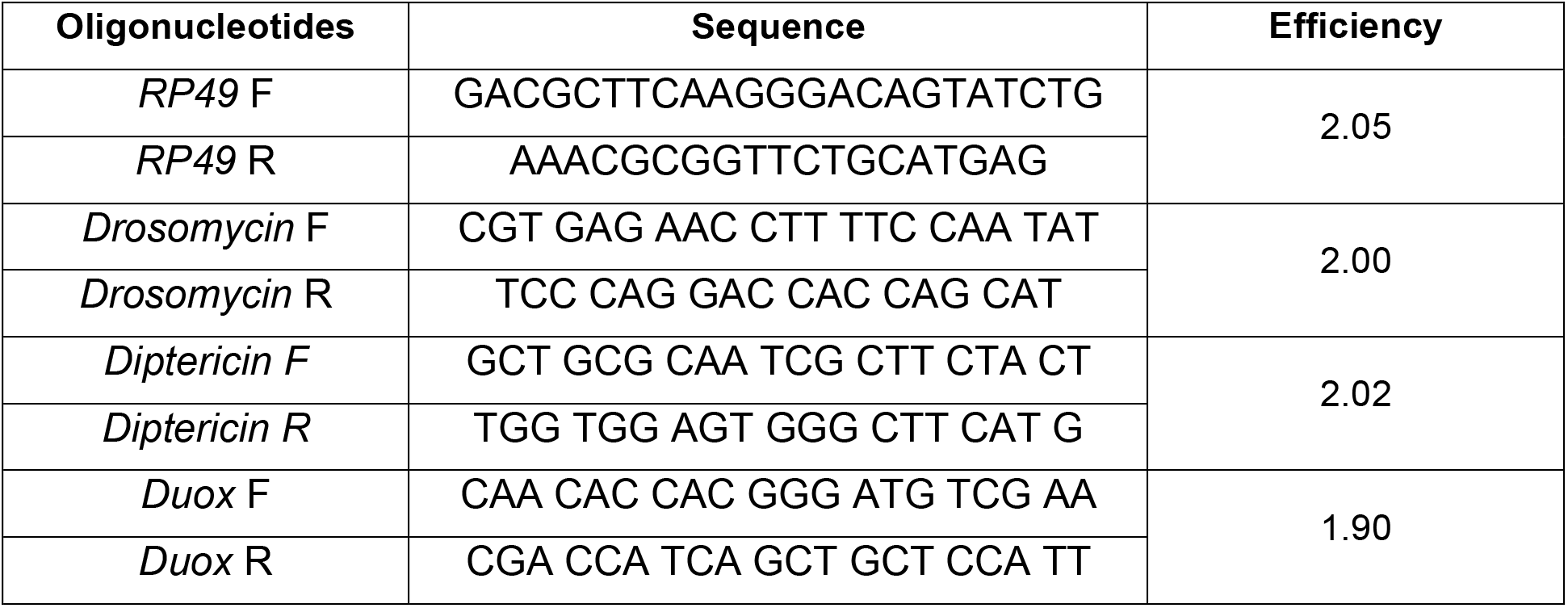

### Statistical analysis

Data are represented as mean ± SEM. Statistical tests were performed with GraphPad (Prism 6). For experiments with two groups of samples, Student’s t-test was performed. For experiment with three groups and more, we applied Two-way ANOVA test. For survival curves, results from 3 independent experiments were grouped (at least 70 animals) and analyzed by Kaplan-Meier test. For Fig S4D, we applied Fisher’s exact test. For qRT-PCRs, at least 15 animals were included per point; otherwise 10 animals were included per point.

**\****P* < 0.05, ** *P* < 0.01, *** *P* < 0.001

## Acknowledgements

We thank D. Ferrandon and S. Liégeois (IBMC, Strasbourg, France), and W.J. Lee (SNU, Seoul, South Korea) for sharing flies stocks and bacteria. We also thank Jost Enninga (Insitut Pasteur, Paris) for the gift of the S. Typhimurium strain expressing DsRed. We are grateful to the core Imaging facility of the “Structure Fédérative de Recherche Necker” INSERM US24 CNRS UMS 3633 for their technical support.

The following reagents were provided by the Network on Antimicrobial Resistance in *Staphylococcus aureus* (NARSA) for distribution by BEI Resources, NIAID, NIH: *Staphylococcus aureus* USA300 WT and *Staphylococcus aureus* USA300 *ΔkatA.* We thank Thierry Capiod (INEM, Paris, France) for technical help.

---

**Figure S1.**
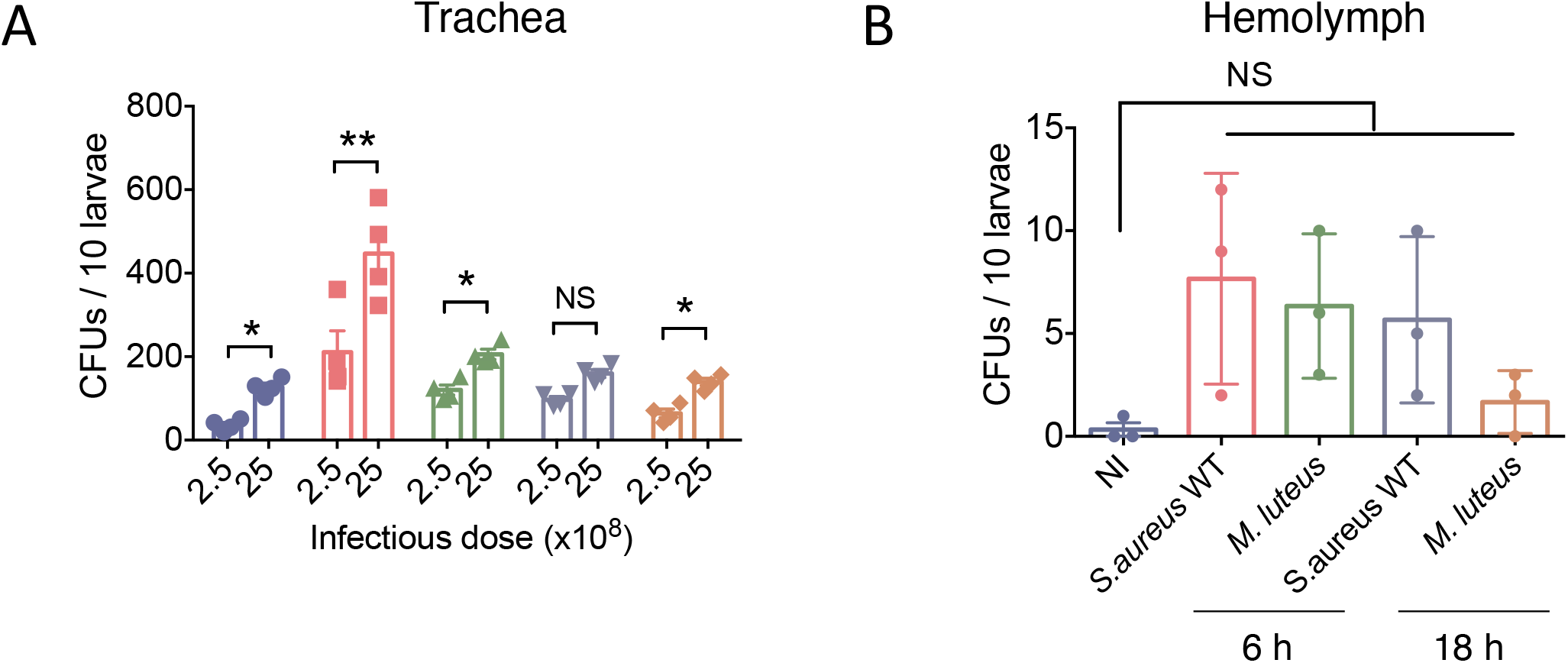
Infection is localized in the intestinal tract. **(A)** *w^1118^ D. melanogaster* larvae were orally infected for 30 min with 2.5×10^8^ and 2.5×10^9^ of chloramphenicol-resistant *S aureus* USA300 WT (carrying pRN11 plasmid). Bacterial counts (CFUs) in the trachea were determined at 6, 12, 18 and 24 h p.i.. Tissues were homogenized in DPBS, serially diluted and plated on BHI agar supplemented with chloramphenicol (10 μg.ml^−1^). Mean ± SEM, n=4, NS = not significant, \**P* < 0.05, \*\**P* < 0.01 using two-way ANOVA. **(B)** *w^1118^ D. melanogaster* larvae were orally infected for 30 min with 2.5×10^9^ of *S aureus* USA300 WT. Bacterial counts (CFUs) in the trachea were determined at 6 and 18 h p.i.. After ethanol washing, animals were bled into DPBS and hemolymph plated on BHI agar. Mean ± SEM, n=3, NS = not significant using one-way ANOVA.

**Figure S2.**
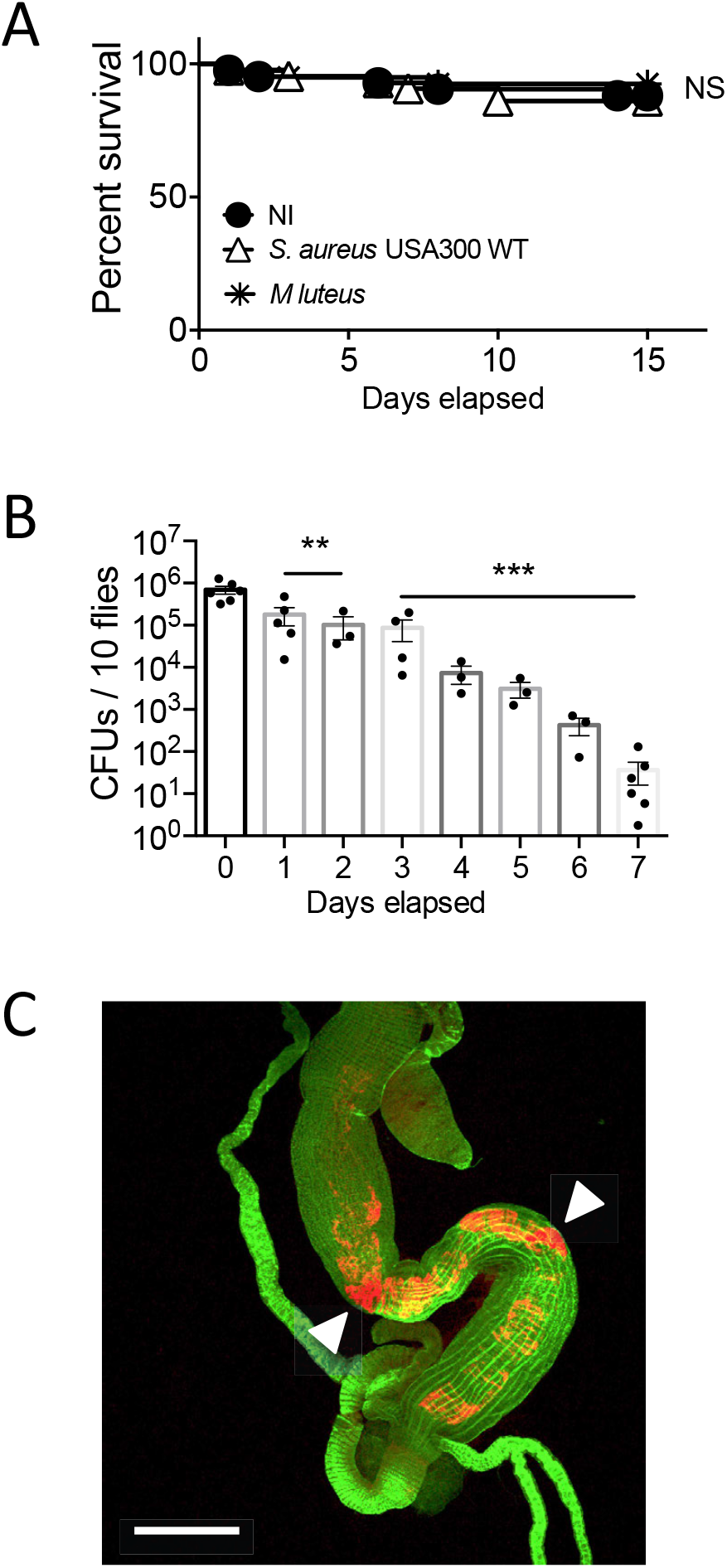
*S. aureus* USA300 is not pathogenic to adults *Drosophila melanogaster.* (**A**) Survival of *w^1118^* adults *D. melanogaster* upon 1 h feeding with an infectious dose of 2.5×10^9^ bacteria of *S aureus* USA300 WT and the non-pathogenic strain *M. luteus*. Animals were followed up every day. 68 animals pooled from 3 independent experiments, NS = not significant using Kaplan-Meier test. (**B**) *w^1118^* adults flies (5 to 7 days old) were orally infected for 1 h with 2.5×10^9^ of chloramphenicol-resistant (CmR) wild-type *S aureus* USA300 bacteria. At indicated time points, guts from 10 flies were homogenized and plated to enumerate intestinal bacteria at the indicated time points. Day 0 corresponds to the end of the proper infection. Mean ± SEM, n=4, \*\**P* < 0.01, \*\*\**P* < 0.001 using one-way ANOVA. (**C**) Representative confocal microscopy imaging of an adult fly intestine, one day after infection with mCherry-*S aureus* USA300 WT. In green, Alexa Fluor™ 488 phalloidin and in red, bacteria expressing the mCherry protein. Scale bar = 0.5 mm. 15 intestines from n=3 and 5 intestines / experiment.

**Figure S3.**
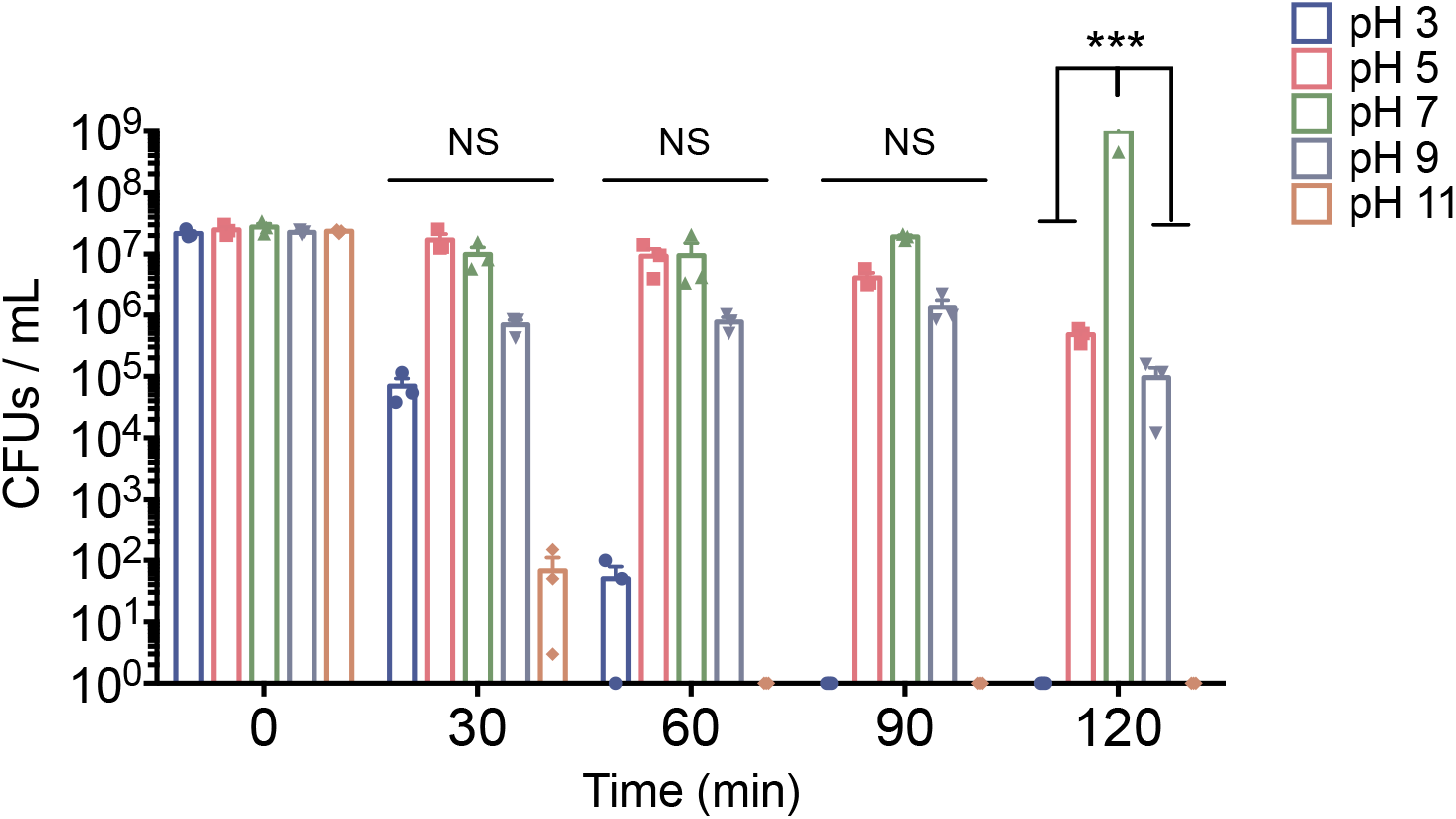
pH tolerance assay of *S. aureus* USA300. Bacteria were inoculated at 2×10^7^ bacteria / mL in BHI broth which pHs were adjusted to 3, 5, 7, 9 or 11. Susceptibility was checked at 30, 60, 90 and 120 min post-inoculation. Mean ± SEM, n=3, NS = not significant, \*\*\**P* < 0.001 using two-way ANOVA.

**Figure S4.**
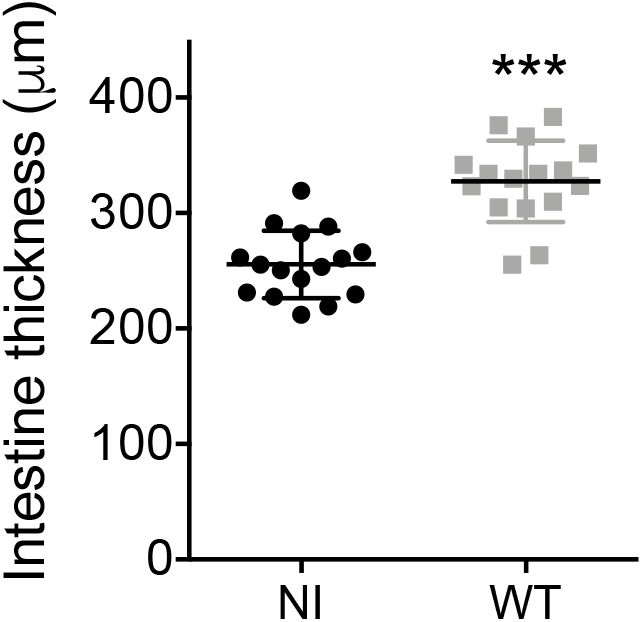
Intestinal inflammation following *S. aureus* USA300 infection. Intestinal thickness measurement of posterior midgut from non-infected (NI) animals and 6 h p.i. with *S. aureus* USA300 (WT). Each point corresponds to one animal. Six animals were dissected for each condition, on 3 independent experiments; data were pooled (n=18). Mean ± SEM, n=3, \*\*\**P* < 0.001 using Mann Whitney test.

**Figure S5.**
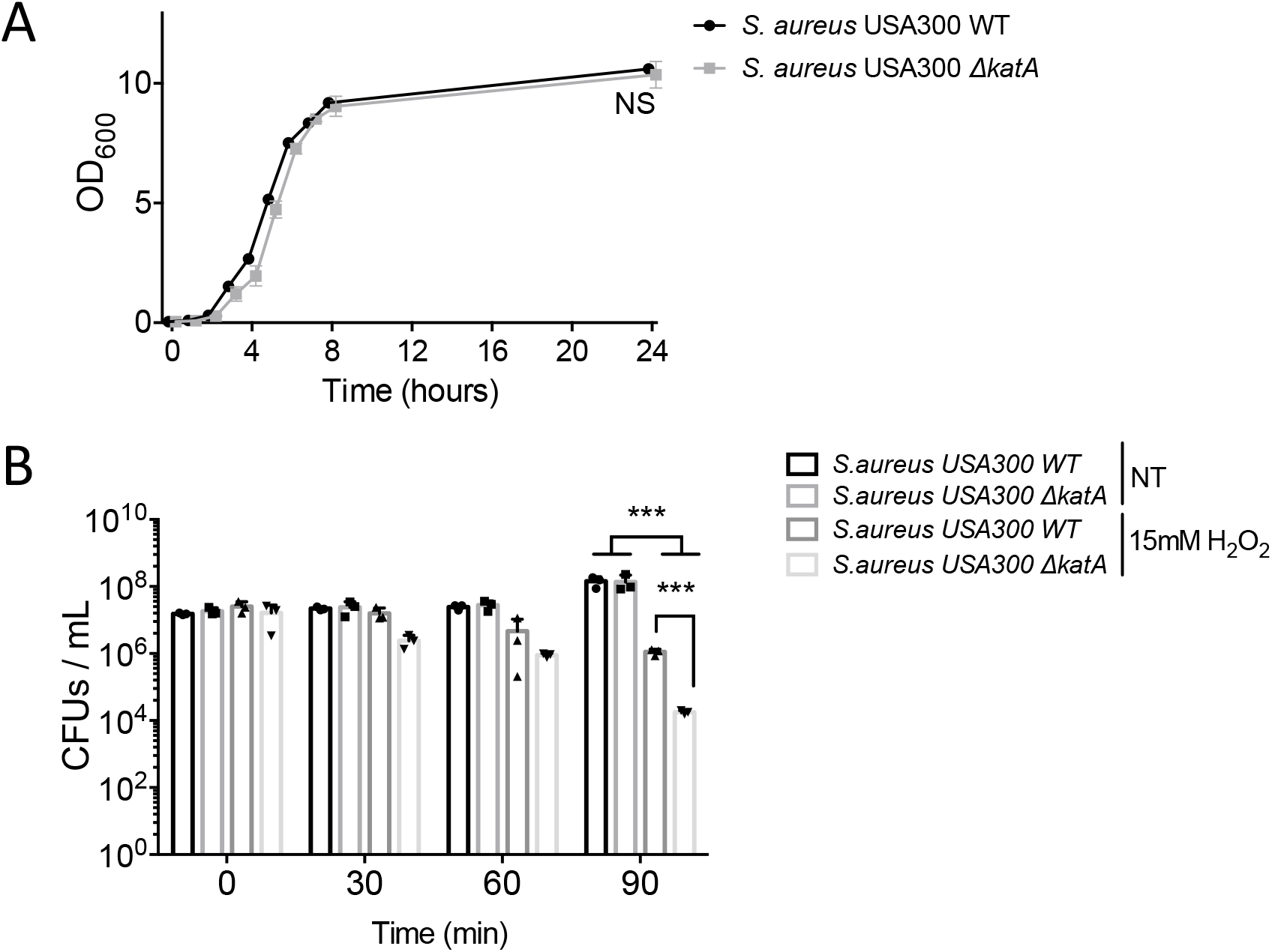
*S. aureus* USA300 *ΔkatA* characterization. **(A)** Growth kinetic of the WT and *ΔkatA* strains in BHI. After overnight growth, bacteria were diluted to a final OD_600_ of 0.05, in 10 mL broth. Every hour, the OD_600_ of the culture was measured, during a 8 h-period and at 24 h. Each point corresponds to the average of 3 experiments. Mean ± SEM, n=3, NS = not significant using one-way ANOVA. **(B)** Exponential phase bacteria were diluted at the concentration 2×10^7^ B / mL and tested for H_2_O_2_ survival in DPBS (15mM). Bacteria were plated each 30 min on BHI agar. Mean ± SEM, n=3, \*\*\**P* < 0.001 using two-way ANOVA.

**Figure S6.**
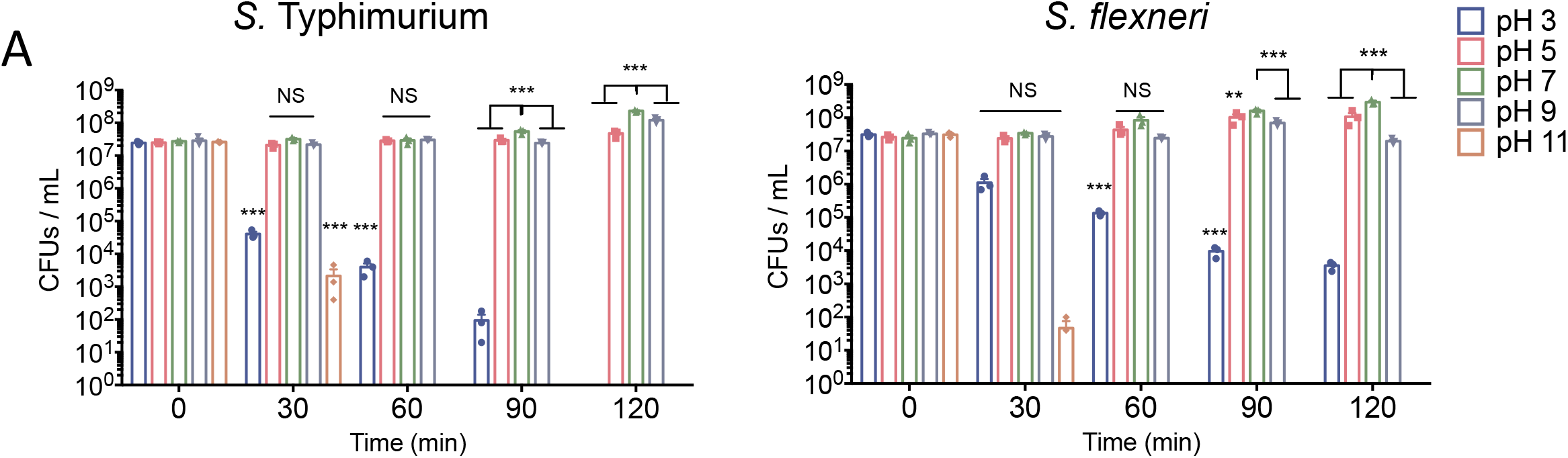
pH tolerance assay of *S. thyphimurium* (left panel) and *S. flexneri (right panel)* Bacteria were inoculated at 2×10^7^ bacteria / mL in BHI broth which pHs were adjusted to 3, 5, 7, 9 or 11. Susceptibility was checked at 30, 60, 90 and 120 min post-inoculation. Mean ± SEM, n=3, NS = not significant, \*\**P* < 0.01 and \*\*\**P* < 0.001 using two-way ANOVA.

**Movie S1, Related to Figure 1 E. Light sheet based imaging of a larvae infected with *S. aureus*.**

Representative posterior mid-half visualization (ventral view) of a mid-L3 larvae, infected with *mCherry* expressing *S. aureus* USA300 WT at the infectious dose 2.5×10^9^ bacteria, for 30 min. At 6 h p.i., animals where fixed and cleared for further Light sheet imaging. Bacteria localize specifically in the intestinal posterior part of the animal.

